# ANU-ADRI and not Genetic Risk score predicts MCI in a cohort of older adults followed for 12 years

**DOI:** 10.1101/070516

**Authors:** Shea J. Andrews, Ranmalee Eramudugolla, Jorge I. Velez, Nicolas Cherbuin, Simon Easteal, Kaarin J. Anstey

**Author notes:** Correspondence to: Shea Andrews, JCSMR, ANU, 131 Garran Rd, Canberra ACT 2601 Australia. Author Email addresses.

## Abstract

**Abstract:** *INTRODUCTION:* We evaluated a risk score comprising lifestyle, medical and demographic factors (ANU-ADRI), and a genetic risk score (GRS) as predictors of Mild Cognitive Impairment (MCI).

*METHODS:* ANU-ADRI risk scores were computed for the baseline assessment of 2,078 participants from the PATH project. Participants were assessed for clinically diagnosed MCI/Dementia and psychometric test-based MCI (MCI-TB) at 12 years of follow-up. Multi-state models estimated the odds of transitioning from cognitively normal (CN) to MCI/Dementia and MCI-TB over 12 years according to baseline ANU-ADRI and GRS.

*RESULTS:* Higher ANU-ADRI score predicted transitioning from CN to either MCI/Dementia and MCI-TB (Hazard ratio [HR] = 1.06, 95% CI:1.04-1.09; HR = 1.06, 95% CI: 1.03-1.09), and a reduced likelihood of cognitive recovery from MCITB to CN (HR = 0.69, 95% CI: 0.49-0.98). GRS was not associated with transition to MCI/Dementia, or MCI-TB.

*DISCUSSION:* The ANU-ADRI may be used for population-level risk assessment and screening.

**Research in Context:** *Systematic Review:* The authors reviewed the literature using online databases e.g. (PubMed). We consulted mild cognitive impairment (MCI) and Alzheimer’s disease (AD) research detailing the use of risk factors for predicting progression from MCI and AD; and the appropriate statistical models for modelling transitions between cognitive states. These publications are appropriately cited.

*Interpretation:* In the general population, the ANU-ADRI comprising lifestyle, medical and demographic factors is predictive of progression from normal cognition to MCI/Dementia whereas a Genetic Risk Score comprising the main Alzheimer’s risk genes is not predictive.

*Future Directions:* Further evaluation of the ANU-ADRI as a predictor of specific MCI and dementia subtypes is required. The ANU-ADRI may be used to identify individuals indicated for risk reduction intervention and to assist clinical management and cognitive health promotion. Genetic risk scores contribute to understanding dementia etiology but apart from *APOE* are unlikely to be useful in screening or prevention trials.

## 1. Background

Accurate risk assessment for cognitive impairment and dementia is increasingly important given the current lack of effective disease modifying treatments for Alzheimer’s disease and other dementias. Risk assessment tools may be used in both pharmacological and non-pharmacological trials, clinics, and for population-level screening to guide risk reduction strategies [1,2]. Validated risk assessment tools that can be administered at very low cost provide methods for low-income countries and regions to assess dementia risk and apply prevention strategies. Given current projections of increasing dementia prevalence, there is an urgent need for validated risk assessment tools that have been evaluated on well characterized samples, over long time periods [3]. However few tools have been evaluated for assessing risk of Mild Cognitive Impairment (MCI) which is a key target group for secondary prevention and pharmaceutical trials.

Recently there has also been an increasing interest in the evaluation of genetic risk scores (GRS) for AD and dementia, which have been associated with the development of AD and incident MCI [4-7], though they have limited utility in predicting AD beyond that attained with basic demographic variables such as age, gender and education [5,8,9]. The number of studies investigating the association AD GRS with progression between cognitive states is limited and the findings mixed. These include reports of a significant association between GRS and progression from to either MCI or Late-onset Alzheimer’s Disease (LOAD) [8] and mixed results for the progression from MCI to LOAD [7,10,11].

Our study has two aims. First, to evaluate the validity of a non-genetic risk score as a predictor of MCI. Our measure [12] is a self-report risk index (the Australian National University Alzheimer’s Disease Risk Index – ANU-ADRI) that has been externally validated in three cohorts of older adults, in which it was found to be predictive of AD and dementia [13]. The second aim is to compare the ANU-ADRI with a GRS. We examine the association between cognitive impairment and the ANU-ADRI and a LOAD GRS, as assessed using a clinical criterion for MCI or dementia and a psychometric test-based criteria for MCI (MCI-TB) in a community-based cohort of older adults using logistic regression and multi-state models.

## 2. Methods

### 2.1 Participants

Participants were community dwelling adults residing in the city of Canberra or the neighboring town of Queanbeyan, recruited into the Personality and Total Health (PATH) Through Life Project, a longitudinal population-based study of health and wellbeing in adults. Cohorts aged 20-24 (20+), 40-44 (40+) and 60-64 (60+) years at baseline were assessed at four-year intervals for a total of 12 years. The background and procedures for the PATH study have been described elsewhere [14]. Written informed consent was obtained from all participants. This study was approved by the Human Research Ethics Committee of The Australian National University.

This study used data from the 60+ cohort, with interviews conducted in 2001-2002 (n = 2,551), 2005-2006 (n = 2,222), 2009-2010 (n = 1,973), and 2014-2015 (n = 1645). Individuals were excluded if they were not Caucasian (n = 107), had a self-reported history of stroke, transient ischemic attack, epilepsy, brain tumours or brain infection 6 (n = 381). As missing values can reduce power and introduce bias in the resulting estimates, missing values that were not attributable to attrition for the predictive variables utilised in the construction of the ANU-ADRI and the test-based MCI (see below) were imputed using an implementation of the Random Forests algorithm available in the ‘missForest’ package in R [15,16]. This left 2,078 individuals available for analysis.

### 2.2 ANU-ADRI risk assessment based on demographic, lifestyle and medical risk factors

The development of the ANU-ADRI and the methodology underlying its computation have been described previously [13]. The ANU-ADRI can be computed based on up to 15 predictive variables, 11 of which are available in PATH, including age (selfreport), gender (self-report), alcohol consumption (calculated according to NHMRC 2001 guidelines [17] using number of drinks per week), education (self reported number of years of education), diabetes (self reported history of diabetes), depression (assessed using the Patient Health Questionnaire (PHQ-9]) [18] following the coding algorithm provided in the PHQ-9 instruction manual with a score of >10 used as cutoff), traumatic brain injury (self reported history of TBI with loss of consciousness), smoking (self reported smoking status for current smoker, past smoker or never smoked), social engagement (constructed from 4 domains for marital status, size of social network, quality of social network, level of social activities. A fifth domain for living arrangements was not available in PATH and thus computed pro rata as the average of the above social engagement variables), physical activity (combined self reported number of hours performing mild, moderate and vigorous activities, weighted by multiples of 1, 2 and 3 respectively [19]), cognitively stimulating activities (assessed as the number of cognitive activities undertaken in the last 6 months for reading, writing, playing games or attending cultural events), and body mass index (BMI equals weight/height^2^, in kilograms/meters^2^). No data were available for the remaining three predictive variables, cholesterol, fish intake and pesticide exposure. The ANU-ADRI is still predictive of the development of dementia even when a subset of variables is used [13]. Values for predictive variables included in the ANU-ADRI for PATH were selected from baseline measurements or the first occasion on which the variables were measured. A constant of +13 was added to the ANU-ADRI to change range to (from -13–19 to 0–32) to facilitate interpretation.

### 2.3 Genotyping and Genetic Risk Score

The most significant LOAD risk SNPs from 23 loci [9,20-24] *ABCA7, BIN1, CD2AP, CD33, CLU, CR1, EPHA1, MS4A4A, MS4A4E, MS4A6A, PICALM, HLA-DRB5, PTK2B, SORL1, SLC24A4-RIN3, DSG2, INPP5D, MEF2C, NME8, ZCWPW1, CELF1, FERMT2* and *CASS4* were genotyped using TaqMan OpenArray assays as previously described [25,26] in addition to the two SNPs defining the APOE alleles which were genotyped using TaqMan assays as previously described [27]. Using these LOAD risk SNPs, an explained variance weighted genetic risk score (EV-GRS) [28] was constructed, which is the sum of all the risk alleles across the individual, weighted by minor allele frequency (MAF) and the Odds Ratio associated with LOAD. The EV-GRS is calculated according to the following formula: 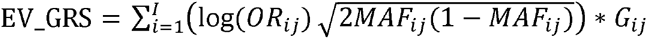 for the *i*th patient, where log(*OR_i_*) the odds ratio for the *j*th SNP; *MAF_ij_* the minor allele frequency for the *j*th SNP; and *G_ij_* = the number of risk alleles for *j*th SNP. Individuals with missing genetic data were excluded (n = 240). We weighted the LOAD SNPs using the previously reported OR for LOAD and by the MAF for the CEU reference population (Supplementary Table 1). The EV-GRS was transformed into a z-score.

### 2.4 Screening and Clinical Assessment

The screening and clinical assessment methods at waves 1-3 have been described elsewhere [29,30] and are briefly summarised here. At each wave, the same predetermined cut-off from a battery of cognitive tests were used for inclusion of participants in a sub-study on mild cognitive disorders and dementia. Participants from the full cohort were selected for clinical assessment if they had any of the following: (i) a Mini Mental State Examination (MMSE) [31] score < 25; (ii) a score below the fifth percentile score on immediate or delayed recall of the first list of the California Verbal Learning Test [32]; or (iii) a score below the fifth percentile on two or more of either the Symbol-Digit Modalities Test [33]; Purdue Pegboard with both hands [34]; or Simple Reaction Time [35]. At wave 4, participants were selected for review if (1) MMSE score <25 or ≤2.5 percentile on one or more cognitive test; or (2) previous diagnosis at waves 1-3; or (3) subjective decline ≥25 on Memory and Cognition Questionnaire (MACQ) or (4) Decline in MMSE score≥3 points.

The criteria for the clinical assessment for cognitive impairment at waves 1-3 has been published by our group elsewhere [30]. It involved a Structured Clinical Assessment for Dementia by one of two physicians, a neuropsychological assessment, and the Clinical Dementia Rating Scale [36], which were used to formulate a consensus diagnosis. Due to the large number of participants screened for review at wave 4, diagnosis was based on neurologist review of interview data, neuropsychological assessment data, self-reported general medical history, psychiatric history, instrumental and basic activities of daily life, and informant reported cognitive and behavioural change, functioning and medical history. For complex cases, two physicians formulated a consensus diagnosis. A detailed protocol for the wave 4 screening procedure and flow chart (Supplementary Figure 1) depicting the screening process for participants is presented in the Supplementary Methods. Clinically diagnosed MCI was based on the Petersen criteria at waves 1 and 2 [37], whereas the Winblad criteria [38] were used at wave 3 and 4. Clinically diagnosed dementia was based on the DSM IV criteria [39] at all waves. At wave 4, there were 14 participants who were not interviewed, but were known to have dementia from informant reports and medical records. Due to the small number of individuals classified with dementia, participants with either MCI or Dementia were grouped into a single MCI/Dementia category.

### 2.5 Test-based MCI

To complement the clinical diagnosis of MCI, a broader psychometric Test-based MCI (MCI-TB) classification was applied to the entire PATH sample [40] at each wave based on education-adjusted cognitive performance (Table 1). The PATH sample was first stratified by education (0-12 or 13+ years). Within each of these strata, individuals were classified as MCI-TB if they scored 1.5 standard deviations below the mean on one or more of the psychometric tests used to assess the following cognitive domains: Perceptual speed, measured using the Symbol Digit Modalities Test [33]; episodic memory, assessed using the immediate recall of the first trial of the California Verbal Learning Test (Recall-immediate) [32]; working memory, measured using the Digit Span Backward from the Wechsler Memory Scale [41]; and vocabulary, assessed by the Spot-the-Word Test [42].

**Table 1:** Characteristics for the PATH cohort for Waves 1 to 4

|  | Wave 1 | Wave 2 | Wave 3 | Wave 4 |
| --- | --- | --- | --- | --- |
| | Estimate ± SD | Estimate ± SD | Estimate ± SD | Estimate ± SD |
| n | 2078 | 1798 | 1596 | 1337 |
| Age | 63 ± 1.5 | 67 ± 1.5 | 71 ± 1.5 | 75 ± 1.5 |
| Female – n (%) | 1009 (49) | - | - | - |
| Education | 14 ± 2.7 | - | - | - |
| Immediate Recall | 7.2 ± 2.3 | 7 ± 2.2 | 6.7 ± 2.2 | 5.4 ± 1.9 |
| Digits Backwards | 4.9 ± 2.2 | 5.1 ± 2.2 | 5.1 ± 2.2 | 5.3 ± 2.2 |
| Spot-the-Word | 52 ± 6 | 53 ± 5.3 | 53 ± 5.1 | 54 ± 5 |
| SDMT | 50 ± 9.7 | 50 ± 9.2 | 48 ± 9.2 | 46 ± 9.5 |
| ANU-ADRI | 9.4 ± 5.9 | - | - | - |
| EV-GRS | 1.6 ± 0.42 | - | - | - |
| **Cognitive Status - n (%)** |  |  |  |  |
| MCI | 23 (1.1) | 28 (1.6) | 35 (2.2) | 103 (7.7) |
| Dementia | 0 (0) | 0 (0) | 7 (0.44) | 37 (2.7) |
| MCI-TB | 384 (18) | 373 (21) | 347 (22) | 261 (19) |
| **Attrition - n (%)** |  |  |  |  |
| Death | - | 57 (2.7) | 54 (3) | 94 (5.8) |
| Dropout | - | 280 (13) | 167 (9.3) | 329 (20.6) |

SDMT: Symbol digits modalities test; MCI: Mild cognitive impairment; MCI-TB:
Test-based mild cognitive impairment

### 2.6 Data analysis

All statistical analyses were performed in R version 3.1.2 [43]. Logistic regression models with the ANU-ADRI and EV-GRS included as covariates in the same model were used to examine their association with MCI/Dementia or MCI-TB status at each wave.

Multi-state models (MSMs) were used to examine the association between the ANUADRI and EV-GRS and transitions between cognitive states. MSMs allow the modelling of competing risks and back transitions between states (i.e., recovery). Hidden Markov models can be used to estimate misclassification error and the effects of covariates can be allowed to vary by transition. A detailed description of multistate-models is provided in the Supplementary Methods. The MSMs utilised in this analysis modelled cognitive deterioration and cognitive recovery by allowing transitions and back transitions between cognitively normal (CN) and cognitively impaired (MCI/Dementia or MCI-TB) states, while ‘death’ was used as a third absorbing state (Figure 1). Individuals with only a single observation (i.e., no recorded transitions) were excluded from the analysis (n = 204). Individuals lost to attrition were considered right censored. The ANU-ADRI and the EV-GRS were included as covariates in the same model. Maximum likelihood estimates of parameters in the MSMs were obtained with the Broyden-Fletcher-Goldfarb-Shanno (BFGS) optimisation method. Normalisation was applied to the likelihood function to improve numerical stability. As the likelihood is maximised using numerical methods, an input of initial values is required to start the search for a maximum. MSMs were fitted using ‘msm’ [44] in R and multiple models were run using different sets of initial values to ensure the robustness of the parameter estimates.

**Figure 1:**
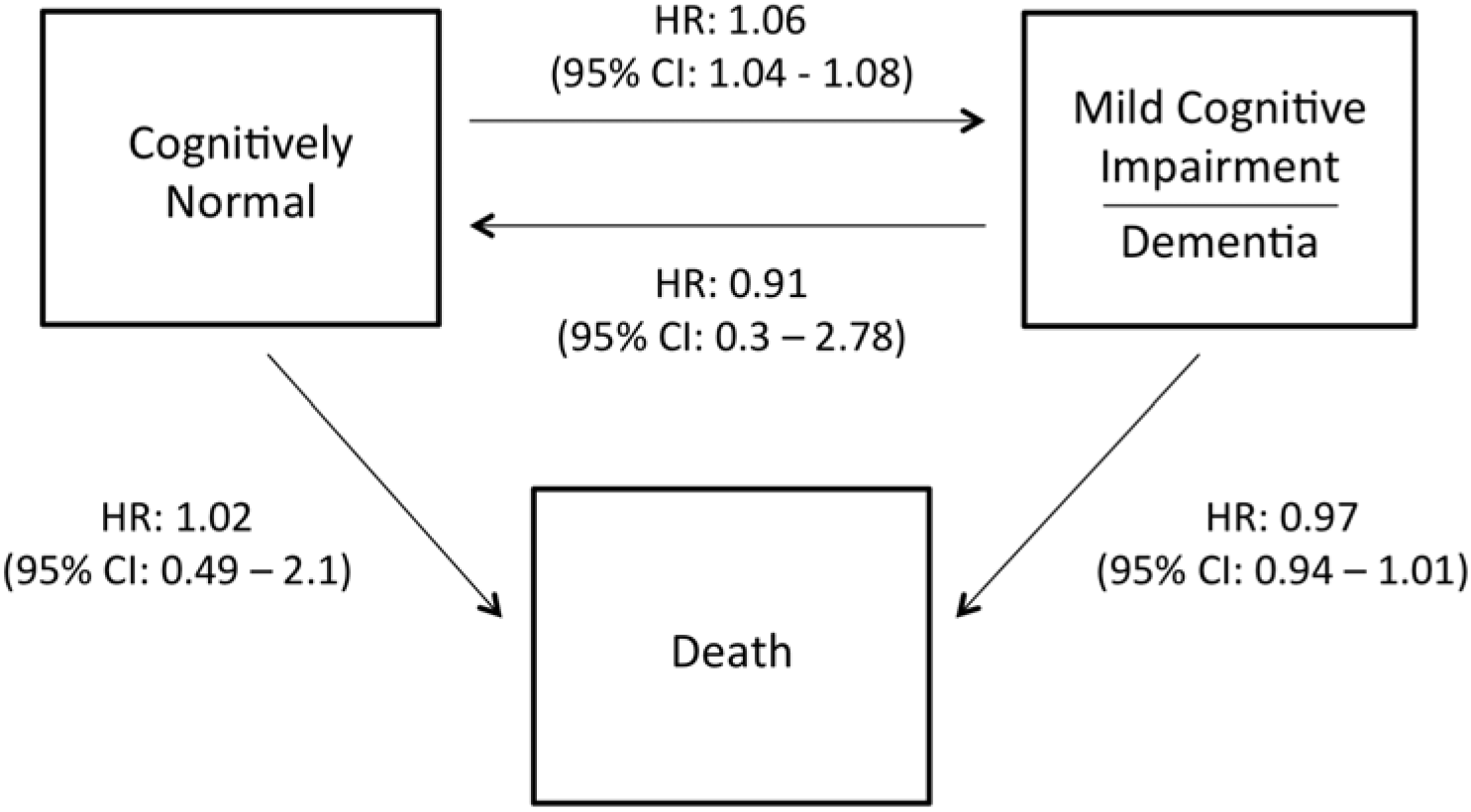
A three state model for possible transitions between cognitive states and death. Hazard ratios (95% confidence intervals) for the effect of the ANU-ADRI on transitions between Cognitively Normal (CN), MCI/Dementia and death are shown. All estimates are from models adjusting for the EV-GRS.

As a sensitivity analysis for the clinical MCI diagnosis, individuals who developed dementia were excluded from the analysis. Additionally, a more stringent criteria MCI-TB was investigated with MCI-TB based on a score of 1.5 SD below the mean on two or more of the above psychometric tests.

## 3. Results

### 3.1 Demographics and other characteristics of the sample

Baseline distributions of education, depression, sex, the ANU-ADRI, raw cognitive tests scores and cognitive states at each wave for the PATH cohort are described in Table 1. Group differences in the sub-indices of the ANU-ADRI between CN and either MCI/Dementia or MCI-TB can be found in Supplementary Table 2. The distribution of the ANU-ADRI scores is shown in Supplementary Figure 2. As expected, the proportion of individuals classified as MCI/Dementia increased over the course of the study, while the proportion of individuals classified as MCI-TB remained stable (Table 1). By wave 4, 36% of the cohort had been lost to follow-up, with 57, 54 and 94 individuals deceased by waves 2, 3 and 4 respectively, and an additional 280, 267 and 329 individuals being lost to follow-up for other reasons (refusal, left catchment area, etc) at waves 2, 3 and 4 respectively.

**Table 2:** Number of transitions between CN, MCI/Dementia and MCI-TB during the length of the study.

| From | To |  |  |  |
| --- | --- | --- | --- | --- |
|  | Cognitively Normal | Impaired | Death | Censored |
| **MCI/Dementia** |  |  |  |  |
| CN | 4460 (86%) | 169 (3.3%) | 189 (3.7%) | 360 (7%) |
| Impaired | 36 (44%) | 37 (41%) | 7 (7.8%) | 6 (6.7%) |
| Censored | 36 (19%) | 3 (1.6%) | 9 (4.8%) | 138 (74%) |
| **MCI-TB** |  |  |  |  |
| CN | 3404 (80%) | 446 (11%) | 144 (3.4%) | 250 (5.9%) |
| Impaired | 321 (31%) | 524 (51%) | 52 (5.1%) | 127 (12%) |
| Censored | 28 (15%) | 11 (5.9%) | 9 (4.8%) | 138 (74%) |

CN: Cognitively normal; MCI: Mild cognitive impairment; MCI-TB: Test-based mild cognitive impairment

Between any two waves, a greater proportion of people transitioned from unimpaired to MCI-TB (11%) than from unimpaired to MCI/Dementia (3%), indicating that MCITB is a more inclusive categorization of cognitive impairment. A smaller proportion of individuals transitioned in the opposite direction - from MCI-TB to unimpaired (31%) than from either MCI/Dementia to unimpaired (44%), indicating that MCI-TB is also a more stable category (Table 2).

### 3.1 Logistic Regression of concurrent and incident MCI

A higher ANU-ADRI (indicating greater risk) score was associated with a higher odds of classification of MCI-TB at baseline and with MCI/Dementia and MCI-TB classifications of cognitive impairment at waves 2, 3 and 4 (Table 3). The EV-GRS was only associated with classification of MCI/Dementia at wave 4.

**Table 3:** Associations between the ANU-ADRI and EV-GRS risk scores and cognitive impairment at waves, 1, 2, 3 and 4. Odds Ratio (95% CI)

| Wave | MCI/Dementia |  | MCI-TB |  |
| --- | --- | --- | --- | --- |
|  | ANU-ADRI<sup>†</sup> | EV-GRS<sup>‡</sup> | ANU-ADRI<sup>†</sup> | EV-GRS<sup>‡</sup> |
| **1** | 1.03 (0.96-1.11) | 0.93 (0.56-1.44) | 1.05 (1.03-1.07)*** | 0.96 (0.85-1.09) |
| **2** | 1.09 (1.03-1.16)*** | 0.76 (0.49-1.15) | 1.07 (1.04-1.09)*** | 1.00 (0.88-1.13) |
| **3** | 1.07 (1.02-1.13)** | 1.03 (0.74-1.42) | 1.05 (1.02-1.07)*** | 1.11 (0.98-1.26) |
| **4** | 1.07 (1.04-1.11)*** | 1.30 (1.08-1.56)*** | 1.05 (1.03-1.08)*** | 1.09 (0.94-1.26) |

\*p < .05; \*\*p < .01; \*\*\*p < .001; MCI/Dementia: Mild cognitive impairment or Dementia; MCI-TB: Test-based mild cognitive impairment; <sup>†</sup> per unitary increase in the ANU-ADRI; <sup>‡</sup> per SD increase in EV-GRS; all estimates are from models adjusting for the ANU-ADRI and EV-GRS

The MCI-TB sensitivity analysis (scoring 1.5 SD below the mean on two or more test) confirmed that the ANU-ADRI was associated with a higher odds of MCI-TB classification at all four waves (OR, *p*; W1: 1.11 [1.07-1.15], <0.001; W2: 1.11 [1.07-1.15], <0.001; W3 1.07 [1.03-1.1], <0.001; W4: 1.08 [1.03-1.12], <0.001). The EVGRS was associated with a reduced odds of MCI-TB classification at wave 1 (OR: 0.7 [0.54-0.9], *p* = 0.007). Excluding dementia cases in the MCI/Dementia sensitivity analysis confirmed that the ANU-ADRI was associated with a higher odds of developing MCI at waves 2, 3 and 4 (OR, *p*: W2: 1.09 [1.02-1.17], 0.01; W3: 1.08 [1.02-1.15], 0.01; W4: 1.07 [1.03-1.11]), <0.001). The EV-GRS was associated with a reduced odds of developing MCI at wave 2 (OR: 0.49 [0.28-0.83], *p* = 0.01).

### 3.3 Multi-state models of transitions

A higher ANU-ADRI score was associated with an increased risk of transitioning from CN to MCI/Dementia and MCI-TB and was negatively associated with cognitive recovery from MCI-TB to CN (Figure 1; Table 4). The probability of transitioning from CN to cognitive impairment after 12 years for individuals scoring 1 SD below the mean on the ANU-ADRI was 11 % and 12.1% for MCI/Dementia and MCI-TB respectively; and for individuals scoring 1 SD above the mean was 22% and 24.7% for MCI/Dementia and MCI-TB respectively.

**Table 4:** Hazard ratios (95% CI) of the ANU-ADRI and EV-GRS scores upon cognitive transition

| Transition | MCI/Dementia |  | MCI-TB |  |
| --- | --- | --- | --- | --- |
|  | ANU-ADRI<sup>†</sup> | EV-GRS<sup>‡</sup> | ANU-ADRI<sup>†</sup> | EV-GRS<sup>‡</sup> |
| **CN - Impaired** | **1.06 (1.04 - 1.08)\*** | 1.08 (0.95 - 1.22) | **1.06 (1.03 - 1.09)\*** | 1.06 (0.85 - 1.31) |
| **CN - Death** | 1.02 (0.49 - 2.1) | 0.7 (0 - 5396.22) | 1.02 (0.96 - 1.07) | 0.82 (0.42 - 1.6) |
| **Impaired - CN** | 0.91 (0.3 - 2.78) | 0.85 (0.03 - 24.35) | **0.69 (0.49 - 0.98)\*** | 0.45 (0.14 - 1.41) |
| **Impaired - Death** | 0.97 (0.94 - 1.01) | 0.89 (0.74 - 1.06) | 1.01 (0.96 - 1.08) | 1.1 (0.59 - 2.04) |

\*p < .05; . CN: Cognitively normal; MCI/Dementia: Mild cognitive impairment or Dementia; MCI-TB: Test-based mild cognitive impairment;
<sup>†</sup>per unitary increase in the ANU-ADRI; <sup>‡</sup>per SD increase in EV-GRS; all estimates are from models adjusting for the ANU-ADRI and EV-GRS

A higher ANU-ADRI score was not associated with transitions from CN, MCI/Dementia or MCI-TB to death; or with cognitive recovery from MCI/Dementia to CN.

The EV-GRS was not associated with either cognitive deterioration or cognitive recovery.

The MCI-TB sensitivity analysis (scoring 1.5 SD below the mean on two or more test) confirmed that the ANU-ADRI was associated with an increased risk of transitioning from CN-MCI-TB (HR: 1.11 [1.06-1.16]), though the negative associated with cognitive recovery was not observed (MCI-TB - CN HR: 1.24 [0.94-1.623]). Excluding dementia cases in the MCI/Dementia sensitivity analysis confirmed that the ANU-ADRI was associated with an increased risk of transitioning from CN-MCI (HR: 1.06, [1.04-1.09]). Additionally, the higher EV-GRS was associated with an increased risk of transitioning from CN-Death (HR: 3.9 [1.3-11.2]).

## 4. Discussion

To our knowledge, no previous study has evaluated both a non-genetic and genetic risk score over a long time period in a population-based cohort. As such this study provides much needed information on the utility of risk assessment approaches for predicting MCI in the general population. Using logistic regression, we found that a per SD increase in the ANU-ADRI score at baseline was associated with a 48-64% and 23-43% increased risk of classification of concurrent or incident MCI/Dementia and MCI-TB respectively, while a per SD increase in EV-GRS was associated with a 30% increased risk of MCI/Dementia classification. Analysis using MSMs indicated that a unitary change in the ANU-ADRI scores is also associated with a 1.06 risk of transitioning from CN to cognitive impairment, and for MCI-TB, 0.69 times reduced risk of cognitive recovery. In contrast, the EV-GRS was not associated with transitions from CN to cognitive impairment or cognitive recovery.

MSMs are well suited to analysing a more ‘realistic’ model of cognitive decline in which cognitive deterioration and recovery are modelled simultaneously in addition to misclassification, death and censoring. This is important in the examination of MCI, as pathological cognitive change is often not a linear progression from normal cognition to MCI and finally to dementia, as reversions from MCI back to normal cognition are common, which was also observed in the PATH cohort [30,45]. Individuals with a stable progression to MCI are more likely to progress to dementia than those with an unstable course or no diagnosis of MCI [45]. A higher ANU-ADRI score is associated both with an increased risk of transition to clinically diagnosed MCI and a reduced chance of reversion back to normal cognition in test-based MCI, suggesting that it could be useful for predicting individuals who are likely to have stable MCI and who are thus at the highest risk of dementia. Additionally, even in individuals who revert to normal cognition, the diagnosis of cognitive impairment may still have prognostic implications as these individuals have a greater likelihood of progressing to dementia or MCI than those who remain cognitively normal [45]. These results show that the ANU-ADRI may be used to measure risk reduction for clinically significant MCI as well as dementia, and have implications for secondary prevention of dementia.

The ANU-ADRI has several strengths [46]. First, the ANU-ADRI is the only risk assessment tool that has not been developed by identifying risk factors through the analysis of a single cohort and as such the predictive variables are not optimised to a particular study. The ANU-ADRI also does not include any risk factors that require clinical assessments or laboratory tests

The findings for the GRS were inconsistent, and difficult to interpret. Across all analysis, there was only one result were the GRS predicted MCI and that was at wave 4, and in the sensitivity analysis, the GRS was actually protective. This lack of an association may be a result of the broad categorization of MCI, rather than MCI subtypes, such that it would have included participants with cognitive impairment that was not MCI due to AD [47,48]. This may also explain the reduced risk associated with both MCI and MCI-TB in our sensitivity analysis. Unfortunately, due to the small number of participants with MCI in PATH, further subgroup analysis would likely be underpowered to detect an effect. However, it should be noted that most dementia cases are associated with mixed pathologies rather than singular pathologies, suggesting that an AD GRS would be associated with both amnestic and non-amnestic MCI [49].

Previous studies have investigated the association of AD GRS with MCI. In 3605 participants (360 MCI, 191 dementia) an AD GRS composed of APOE + 19 LOAD GWAS variants was associated with an increased risk of incident MCI and nominally associated with amnestic and non-amnestic [7]. In a second study of 2674 participants (347 MCI, 132 LOAD) a GRS composed of APOE + 9 LOAD GWAS variants, was associated with progression from to normal cognition to MCI/LOAD [8]. Lack of replication in this study could be due to younger and fewer cognitively impaired participants.

Limitations of our study include the relatively high level of education of the PATH cohort [14]; the ethnicity in PATH is predominately Caucasian, potentially limiting the generalizability of the results in this study to other ethnicities, and biomarkers of AD were not available (e.g. CSF, A). Not all the predictive variables for the ANUADRI was available in PATH, suggesting the present study may underestimate the sensitivity of this tool in predicting individuals who are at risk of developing cognitive impairment. However, the validation studies also included a subset of the variables contributing to the ANU-ADRI [13].

Study strengths included the large population-based sample with high retention rates and twelve years of follow-up data. The PATH cohort was recruited from a narrow age-band, reducing the impact of age-differences on findings. This is particularly important because age has the largest weighting of risk factors in the ANU-ADRI. Finally, the conservative clinical classifications of MCI/Dementia, based on a thorough clinical assessment and consensus diagnosis by clinicians using published criteria, was complemented by a broader psychometric test-based classification of MCI.

In conclusion, higher ANU-ADRI scores predict incident MCI and lower scores predict recovery from MCI to normal ageing. These results complement previous evidence that the ANU-ADRI is predictive of AD and dementia [13]. In comparison, a genetic risk score comprising the main AD genes did not predict MCI in the population. These results provide further support for using the ANU-ADRI for individual patient assessment and for informing intervention and treatment strategies aimed at delaying or preventing dementia.

## Acknowledgments

We thank the participants of the PATH study, Peter Butterworth, Andrew Mackinnon, Anthony Jorm, Bryan Rodgers, Helen Christensen, Patricia Jacomb and Karen Mawell. The study was supported by the Dementia Collaborative Research Centres, the National Health and Medical Research Council (NHMRC) grants 973302, 179805, and 1002160. 1002560. JIV was supported by the Eccles Scholarship in Medical Sciences, the Fenner Merit Scholarship and the Australian National University High Degree Research scholarships. NC is funded by Research Fellowship No. 12010227. KJA is funded by NHMRC Research Fellowship No. 1002560.

